# Oligosaccharyltransferase is involved in targeting to ER-associated degradation

**DOI:** 10.1101/2024.05.12.593735

**Authors:** Marina Shenkman, Navit Ogen-Shtern, Chaitanya Patel, Bella Groisman, Metsada Pasmanik-Chor, Sonya M. Schermann, Roman Körner, Gerardo Z. Lederkremer

## Abstract

Most membrane and secretory proteins undergo N-glycosylation, catalyzed by oligosaccharyltransferase (OST), a membrane-bound complex in the endoplasmic reticulum (ER). Proteins failing quality control are degraded via ER-associated degradation (ERAD), involving retrotranslocation to cytosolic proteasomes. Using SILAC proteomics, we identified OST subunits as key interactors with a misfolded ER protein bait, suggesting unexpected involvement in ERAD. Previous reports implied additional roles for OST subunits beyond N-glycosylation, such as quality control by ribophorin I. We tested OST engagement in glycoprotein and non-glycosylated protein ERAD; overexpression or partial knockdown of OST subunits interfered with ERAD in conditions that did not affect glycosylation. Effects were studied on misfolded membrane proteins, BACE476 and asialoglycoprotein receptor H2a, and the luminal α1-antitrypsin NHK variant. OST appears to participate in late ERAD stages, interacting with the E3 ligase HRD1 and facilitating retrotranslocation. Molecular dynamics simulations suggest membrane thinning by OST transmembrane domains, possibly assisting retrotranslocation via membrane distortion.

## Introduction

Secretory proteins are translocated into the endoplasmic reticulum (ER), where they undergo folding and quality control. Molecules that are terminally misfolded are retrotranslocated to the cytosol to be polyubiquitinated and degraded by proteasomes, in a process known as ER-associated degradation (ERAD) ^1,2^. The precise mechanism of retrotranslocation remains a subject of ongoing study. It appears that various mechanisms may exist for different ERAD substrates, contingent upon the specific E3 ubiquitin ligase involved. Among these ligases, the most well-characterized participant in ERAD is HRD1, which forms a large protein complex mediating retrotranslocation ^3,4^. Studies using genetic screens in yeast identified ERAD complexes involving HRD1 ^5^. Subsequently, analogous components were found in mammals, including HRD1 itself ^6^. The primary constituents of the mammalian HRD1 complex encompass transmembrane proteins such as Derlin 1-3, Herp, and Ubxd8, along with SEL1L, OS9, and XTP3-B on the luminal side. On the cytosolic side, components include Ubc6 or Ube2G2, p97, and NGLY1 ^3,5^. Structural insight into the HRD1 complex has been reported in yeast but not yet in mammalian cells ^7,8^. Cryo-electron microscopy (CryoEM) studies of the yeast HRD1/HRD3 complex indicated a direct role in retrotranslocation ^7^. Further evidence involves *in vitro* reconstitution experiments suggesting a regulation of retrotranslocation by HRD1 autoubiquitination ^9,10^. Herp, a protein tethered to the ER membrane via a hydrophobic hairpin, with most of the protein on the cytosolic side, was shown to recruit HRD1 and misfolded proteins for ERAD at a specialized ER subcompartment, the ERQC ^11,12^.

An approach used to delineate the HRD1 complex and discover new components has been the pull-down of the complex through known ERAD components, such as Yos9 in yeast ^13^ and SEL1L ^14^, XTP-3B, HRD1, UBXD8, Derlin-1/2 and OS-9 in mammalian cells ^15^. The caveat is that overexpression affects the function and interactions of these proteins. Furthermore, even when tagged by knock-in at the endogenous level the isolated complexes might be non-functional ^16^.

This is because these complexes might be dormant, partial complexes, given the dynamic nature of complex assembly, influenced by the presence of an ERAD substrate ^11,17^. Here we have used a SILAC (stable isotope labeling by amino acids in cell culture) proteomics approach to capture functional ERAD components. We accomplish this by pull-down of an ERAD substrate and analyzing its preferential interactors under conditions favoring the accumulation of molecules targeted for retrotranslocation and subsequent proteasomal degradation. Surprisingly, among the most highly enriched interactors were several subunits of the oligosaccharyltransferase (OST) complex.

OST plays a pivotal role in N-glycosylation, the primary post-translational modification of secretory proteins. It transfers a conserved oligosaccharide from a lipid-linked glycan donor to asparagine residues in defined sequons in newly-synthesized proteins. In mammalian cells, the OST complex consists of a catalytic transmembrane subunit (STT3A or STT3B) along with several additional transmembrane proteins including DAD1, OST48/DDOST, TMEM258, OST4, ribophorin1, ribophorin2, and subunits interacting with STT3A (KCP2, DC2/OSTC) or STT3B (TUSC3/N33 or IAP/MAGT1) ^18,19^. While the primary function of the OST complex lies in protein N-glycosylation, studies suggest the participation of OST subunits, such as ribophorin I, in protein quality control ^20,21^, although the mode of action remains unclear. Here we show that OST subunits have a role in protein targeting to ERAD.

## Results

### Identification of OST as ERAD substrate interactor by a SILAC proteomics approach

The proteomics approach that we devised involved making a construct of an established model ERAD substrate, H2a, with an affinity binding tag (streptavidin binding peptide, SBP) and producing a stable HEK293 cell line (SBP6) expressing this fusion protein, H2aSBP. H2a, the uncleaved precursor of human asialoglycoprotein receptor (ASGPR) H2a, has been extensively studied. H2a is a type II membrane glycoprotein, completely retained in the ER ^22^ and is a substrate of the HRD1 complex ^11,23^. As H2a can undergo natural cleavage, generating a soluble luminal fragment, we used an uncleavable mutant, H2aG78R ^24^. The use of the 38 amino acid SBP affinity tag allows for specific and high affinity binding and elution under mild conditions, to preserve complexes ^25^. H2aSBP behaves like H2a in its ER retention and targeting to ERAD and H2aSBP precipitation with streptavidin beads allows coprecipitation of Derlin1, HRD1 and Fbs2, with the latter interaction being inhibited with kifunensin, which inhibits mannose trimming required for ERAD ^23^.

SBP6 cells were cultured in the presence of media containing heavy amino acids (13C(6)15N(2) lysine) to uniformly label the proteins, while WT HEK293 cells were grown with normal light amino acids, or vice versa (Fig. 1a). Cell lysates in mild non-denaturing detergent (1% digitonin) from cells with or without H2aSBP expression were mixed at a 1:1 protein ratio, followed by precipitation with streptavidin-conjugated beads and elution by competition with biotin. Eluted proteins were subjected to SDS-PAGE, digested with trypsin and identified by electrospray tandem mass spectrometry (ESI-MS/MS). This experiment was done in cells treated without or with proteasomal inhibitors to accumulate the ERAD substrate before degradation. Both cells treated with or without proteasomal inhibitors were incubated briefly with puromycin before cell lysis, to reduce the amount of proteins undergoing cotranslational translocation into the ER. Puromycin causes the termination of nascent polypeptide chains and their release into the lumen of the ER ^26^.

**Fig. 1.**
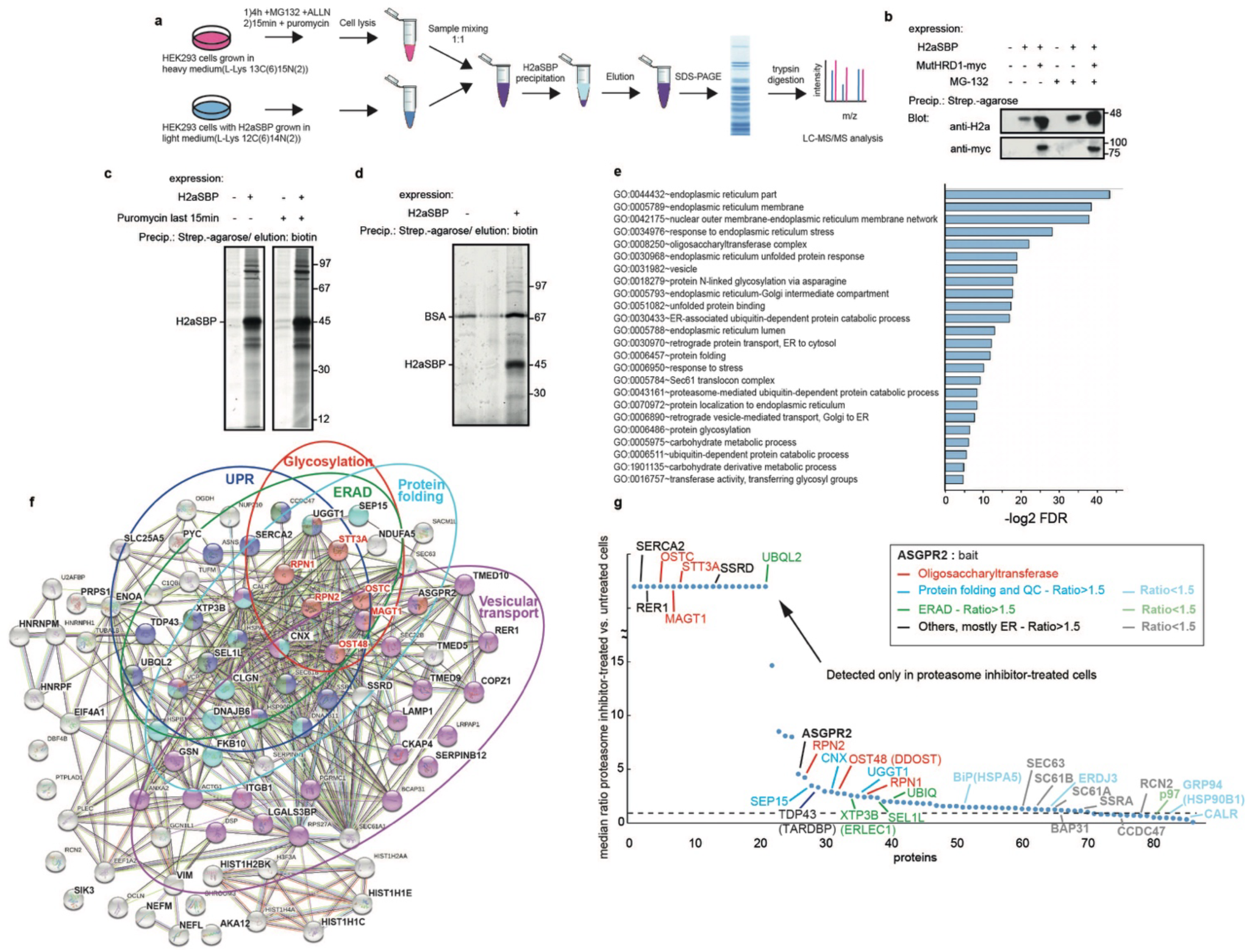
OST subunits are among top ERAD substrate interactors identified by a SILAC proteomics approach. **(a)** Experimental scheme of SILAC-based mass spectrometry analysis of HEK293 cells labeled with lysine containing heavy isotopes for 10 days or HEK293 stably expressing H2aSBP (SBP6 cell line) without heavy lysine (or vice versa). Cells were treated with MG-132 (50μM) and ALLN (50μM) for 4h and puromycin (2mM) for 15 min before lysis, or only with puromycin. Cells were solubilized in 1% digitonin in PBS and lysates from cells labeled with or without heavy lysine were mixed 1:1. Protein complexes pulled down with H2aSBP (using streptavidin-agarose beads) were eluted with 250μM biotin, subjected to SDS-PAGE followed by trypsin digestion and LC-MS/MS analysis. **(b)** Effect of mutant HRD1 on H2aSBP. Dominant negative mutant myc-tagged hsHRD1 (mutHRD1-myc) was transfected into SBP6 cells. Two days post-transfection these cells or control HEK293 cells were treated or not for 4h with 20 μM MG132, lysed and subjected to precipitation with streptavidin beads. Biotin-eluted samples were subjected to 10% SDS-PAGE and immunoblotted with anti-H2a or anti-myc. MW markers in kDa are on the right. **(c)** Metabolically labeled proteins pulled down with H2aSBP. SBP6 cells or control HEK293 cells were labeled with [^35^S] Cys/Met mix for 5h under proteasomal inhibition with 20mM MG132 and treated or not with 2mM puromycin for the last 15 min before lysis. Cells were lysed in digitonin buffer followed by precipitation with streptavidin beads, elution with biotin and SDS-PAGE with detection by fluorography. **(d)** Proteins pulled down with H2aSBP and silver stained. SBP6 cells or control HEK293 cells were treated for 4h with 20μM MG132 and 2mM puromycin (last 15 min), lysed in digitonin buffer followed by precipitation with streptavidin beads (in the presence of 0.1% BSA), elution with biotin, SDS-PAGE and silver staining. **(e)** Gene Ontology (GO) analysis of differentially expressed genes from the SILAC results of cells with or without proteasomal inhibition (ratio>1.5). -log2 enriched p-values are shown for selected top entries with the lowest false discovery rates. (see also Suppl. Table 1). **(f)** STRING interaction map of proteins identified in the SILAC analysis, highlighting those with >1.5 ratio when comparing cells treated with or without proteasomal inhibitors. OST protein subunits are in red, other enriched functions are in colored circles. **(g)** Ratio of levels of proteins identified in cells treated with or without proteasomal inhibitors. The proteins on the left (up to UBQL2) were detected only in proteasomal inhibitor-treated cells. The dotted line is a value of 1, equal presence in cells treated with or without proteasomal inhibitors. (see also Suppl. Tables 2 and 3).

The expression of H2aSBP increased by proteasomal inhibition and more robustly by overexpression of a dominant negative RING finger mutant of HRD1, as expected for an HRD1 substrate. The mutant HRD1 coprecipitated with H2aSBP upon pull-down of the latter with streptavidin-agarose beads (Fig. 1b).

The SBP tag allowed specific and efficient elution in non-denaturing conditions with a competitive concentration of free biotin, showing many proteins coprecipitated specifically from a lysate of cells labeled with [^35^S] met+cys in the presence of the proteasome inhibitor MG-132 (Fig. 1c). Brief treatment with puromycin before lysis did not decrease the signal of these proteins. We used mild lysis conditions (see Methods), combined with the gentle biotin elution, to enable preservation of protein complexes with little or no non-specific precipitation. Silver staining of the eluates after SDS-PAGE in a similar experiment showed an excellent signal to noise ratio, with only very weak bands in the control without H2aSBP, except for BSA, which was used as a blocker to reduce non-specific binding during the affinity pull-down. (Fig. 1d).

After applying the SILAC proteomics protocol (Fig. 1a), we selected as interactors only hits showing >2 fold ratio of labeled/ unlabeled after normalization to the total amount of H2a-SBP (bait) obtained. We then compared the results from cells treated for 4h with MG-132 and ALLN and 15 min with puromycin to those from cells treated only with puromycin. Gene ontology analysis showed that proteasomal inhibition led to a significant enrichment for cellular functions related to the ER, ER membrane and ubiquitin/proteasome processes, which were expected, and gave us confidence in the approach. One of the top function enriched categories using the DAVID gene enrichment tool was the OST complex (p-value=3.13e^-10^), which was unexpected (Fig. 1e). A STRING interaction map showed high enrichment of proteins belonging to a few categories related to protein quality control: ERAD (p-value=1.28e^-7^), protein folding (p-value=1.28e^-7^), glycosylation (p-value=7.03e^-10^)(including OST and calnexin (CNX) cycle genes), unfolded protein response (UPR) and vesicular transport (p-value=8.00e^-6^)(Fig. 1f). This latter category perhaps relates to the participation of vesicular trafficking in ERAD, which we reported recently ^27^.

Among the most significant H2aSBP interactors, with enrichment values of 2-fold or higher in proteasomal inhibitor-treated cells, were most OST subunits (Table 1). This was even more surprising, considering that membrane proteins, some with multiple membrane spans such as OST subunits, are usually underrepresented in the MS analysis. All of these OST subunits were highly enriched upon proteasomal inhibition (Fig. 1g). It should be stressed that the puromycin treatment before cell lysis should have reduced interactions with newly-made translocated proteins, where H2aSBP could have encountered OST, which is associated with forward translocons ^28^. Indeed, levels of forward translocon related proteins, such as Sec61 subunits or SSRA were at similar or reduced levels in samples treated with or without proteasomal inhibitor (Fig. 1g). Other enriched proteins in proteasomal inhibitor-treated samples include ERAD-related genes, such as XTP3-B, SEL1 and ubiquitin, calnexin cycle genes (CNX, SEP15 and UGGT1) and other relevant genes such as the ER calcium pump SERCA2 and TDP43 (Fig. 1g). TDP43, normally localized in the nucleus, accumulates in the cytoplasm in several diseases, including ALS, and also upon proteasome inhibition ^29^.

**Table 1.** OST subunits detected by mass spectrometry. LC-MS/MS analysis following the experimental scheme of Fig. 1a. Results of >2 fold labeled/unlabeled from cells treated with MG-132 and ALLN for 4h and puromycin for 15 min before lysis were compared to those from cells treated with only puromycin for 15 min, after normalization to the total amount of H2a-SBP (ASGPR2, bait) obtained. Hits are ranked by enrichment in cells treated with vs. without proteasome inhibitors. Only OST subunits, with enrichment values of 2-fold or higher in cells treated with vs. without proteasome inhibitors or found only in cells treated with proteasome inhibitors are shown. See Suppl. Tables 2 and 3 for complete lists of hits.

| OST subunit | UNIPROT Accession No. | Hit No. | Median ratio with vs. without proteasome inhibition | Peptides (seq) | Sequence Coverage (%) |
| --- | --- | --- | --- | --- | --- |
| IAP/ MAGT1, MAGT1_HUMAN | Q9H0U3 |  | Only in cells treated with proteasome inhibitors | 4 | 10.1 |
| STT3 A, STT3A_HUMAN | P46977 |  | Only in cells treated with proteasome inhibitors | 8 | 11.2 |
| DC2/ OSTC, OSATC_HUMAN | Q9NRP0 |  | Only in cells treated with proteasome inhibitors | 1 | 8.1 |
| Ribophorin II, RPN2_HUMAN | P04844 | 27 | 4.3 | 15 | 30.9 |
| OST 48, OST48_HUMAN | P39656 | 32 | 2.8 | 11 | 29.8 |
| Ribophorin I, RPN1_HUMAN | P04843 | 36 | 2.4 | 18 | 32.5 |

### Colocalization and increased interaction of an OST subunit with the ERAD substrate upon proteasomal inhibition

Given the increase in the pull-down of OST subunits with the ERAD substrate upon proteasomal inhibition, we then analyzed whether there was any change in the subcellular localization of an OST subunit in relation to the ERAD substrate under these conditions. We chose to examine the OST subunit Defender Against apoptotic cell Death 1 (DAD1) ^30^ because it was well characterized in previous studies, although it was not detected in our proteomics analysis, possibly due to its low abundance. GFP-DAD1 had been shown to associate with forward translocons ^31^. We compared the localization of GFP-DAD1 to the ERAD substrate H2a linked to monomeric red fluorescent protein (H2a-RFP). Both GFP-DAD1 and H2a-RFP show an ER pattern in untreated cells, although with some concentration in the juxtanuclear region (Fig. 2a). Surprisingly, under proteasomal inhibition with lactacystin (Lac), GFP-DAD1 accumulated together with H2a-RFP in the juxtanuclear ER-derived quality control compartment (ERQC), a staging ground for ERAD ^12,24,32^, (Fig. 2b). This is similar to the effect of ERAD substrate accumulation (upon proteasomal inhibition or by other means) on the localization of ERAD machinery components (HRD1, Derlin-1 ^23^). Several ER chaperones, such as CNX and CRT, also relocalize to the ERQC under these conditions, but not others, such as BiP ^32,33^. As we had also seen before ^24^, the translocon subunit Sec61β linked to GFP accumulates at the ERQC with the ERAD substrate upon proteasomal inhibition (Fig. 2c, d). To test whether the entire translocation complex linked to ribosomes (at the rough ER) accumulates in the ERQC along with the ERAD substrate, we immunostained cells transfected with H2a-RFP using an anti-ribosomal RNA (rRNA) antibody as a ribosomal marker. As opposed to DAD1 and Sec61β, the ribosomes did not accumulate together with H2a-RFP after Lac treatment and remained in a cytosolic and ER pattern (Fig. 2e, f). Sec61 participates in the forward translocation complex, but has also been implicated in retrotranslocation ^34^.

**Fig. 2.**
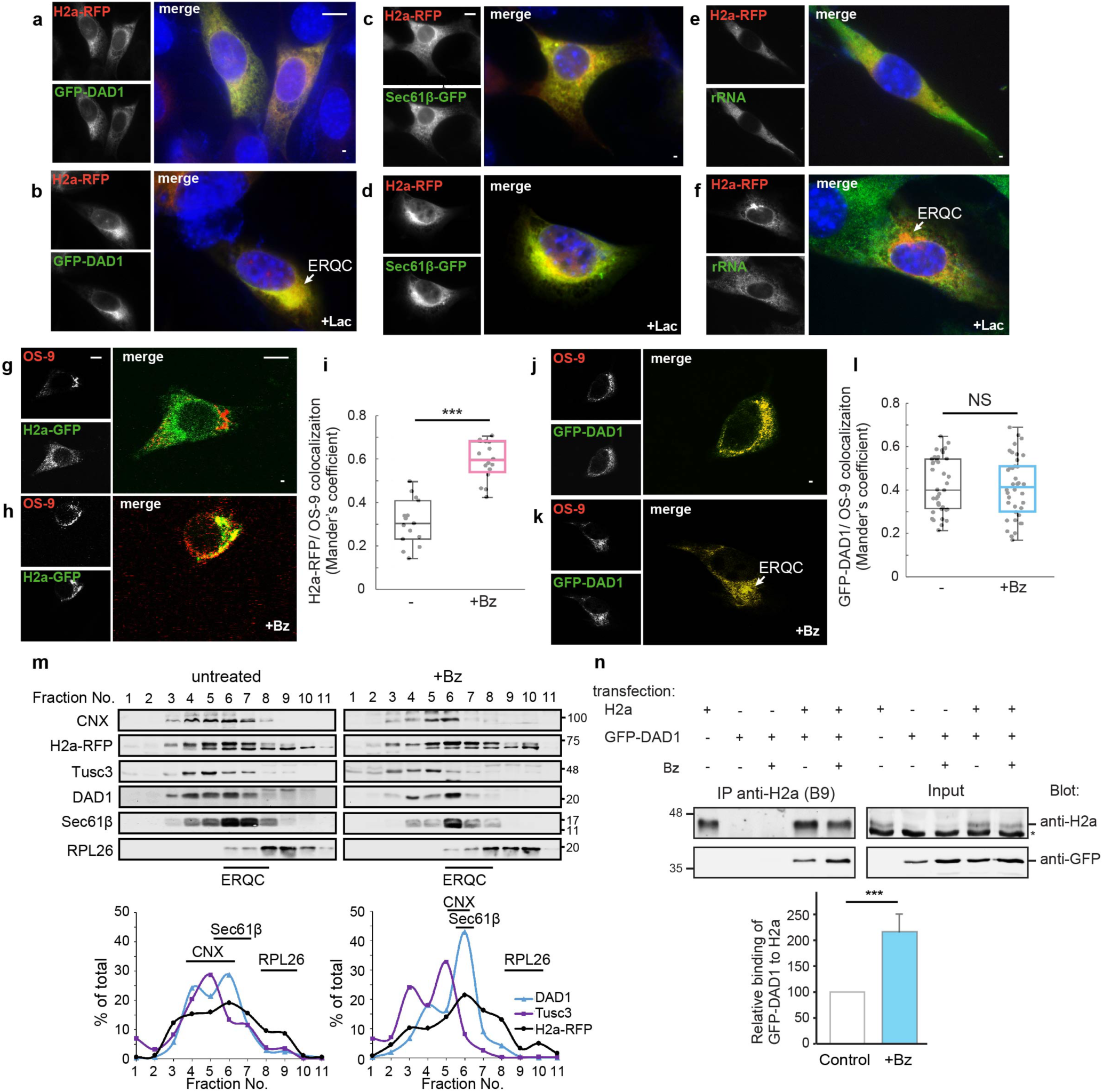
DAD1 accumulates in the ERQC upon proteasomal inhibition separately from ribosomes and interacts increasingly with an ERAD substrate. **(a-d)** GFP-DAD1 and Sec61β-GFP accumulate at the ERQC. NIH 3T3 cells were transiently cotransfected with the ERAD substrate H2a-RFP and with GFP-DAD1 or Sec61β-GFP and were incubated with or without 17 μM Lac for 3h. **(e-f)** Ribosomes do not accumulate at the ERQC. NIH 3T3 cells transiently transfected with H2a-RFP were treated as in (a-d) and were fixed and immunostained using a primary mouse anti-rRNA antibody and secondary goat anti-mouse IgG antibody conjugated to Cy2. **(g-l)** GFP-DAD1 colocalizes with OS-9 with or without proteasomal inhibition. S-tagged OS-9.1/2 was expressed in NIH 3T3 cells together with H2a-GFP or GFP-DAD1, with or without treatment with 1.5 μM Bz for 3h, followed by fixation and immunostaining with mouse anti-S tag antibody and secondary goat anti-mouse IgG antibody conjugated to Dylight 549. The graphs are from data from 3 repeat experiments. **(m)** Comigration of endogenous DAD1 and the ERAD substrate at the ERQC. HEK293 cells transiently transfected with H2a-RFP were incubated without (left) or with Bz (2 μM) for 3 h. Cells were homogenized and postnuclear supernatants were centrifuged on 10–34% iodixanol gradients. Fractions of the gradients were run on SDS–PAGE and immunoblotted with the indicated antibodies. H2a-RFP was detected with anti-RFP. A representative experiment out of 3 repeat experiments is shown. **(n)** Immunoprecipitation with a mouse monoclonal (B9) anti-H2a of lysates of HEK293 cells expressing H2a together with GFP-DAD1, with or without treatment with 2 μM Bz for 3 h. Crosslinking with 1 mM DSP was performed before cell lysis. Immunoprecipitates and 10% of the total lysates (input) were immunoblotted with rabbit anti-H2a amino-terminal antibody and anti-GFP. Asterisk= non-specific band. The graph shows increased coimmunoprecipitation of GFP-DAD1 with H2a upon proteasomal inhibition. Values are relative to control (=100), normalized by total GFP-DAD1 and H2a, average of 4 independent experiments ±SD. p value = 0.0003.

We compared the localization of GFP-DAD1 with that of the ERAD-associated luminal lectin OS-9, which appears constitutively localized mainly at the ERQC ^11,35^. Similar to H2a-RFP, the GFP-DAD1 that accumulated under proteasomal inhibition colocalized with OS-9 (Fig. 2 h, k). In this experiment bortezomib (Bz) was used for proteasome inhibition. However, while the colocalization of H2a-RFP with OS-9 increased significantly upon proteasomal inhibition, that of GFP-DAD1 did not change (Fig. 2 i, l). This suggests that a portion of GFP-DAD1 molecules localize to the ERQC in untreated cells and may already colocalize with OS-9 in peripheral regions.

We used iodixanol gradients optimized to separate ER subcompartments as a different approach to analyze the localization of these proteins in relation to the ERQC ^11,36^. As we had seen before ^11^, H2a-RFP appeared mainly in middle fractions of the gradient (5-7), with a small shift to heavier fractions (6-8), upon proteasomal inhibition with Bz (Fig. 2m). Endogenous DAD1 showed a similar pattern in untreated cells, with a sharper accumulation in Bz-treated cells, as a peak in fraction 6, and an additional smaller peak in fraction 4. Endogenous CNX and Sec61β also appeared in middle fractions, partially overlapping with DAD1 in untreated conditions and in Bz-treated cells where they concentrated in fraction 6. We analyzed another OST subunit, Tumor suppressor candidate 3 (Tusc3), a non-essential subunit that was also absent from our top proteomic hits. Endogenous Tusc3 showed a similar pattern to DAD1 in untreated conditions, although concentrating more at fraction 4, but it did not shift to higher densities upon proteasomal inhibition (Fig. 2m). A ribosomal subunit, RPL26 appeared at higher densities, marking regions of rough ER, with little or no overlap with the ERQC (Fig. 2m), as we had seen before ^11^.

A specific interaction of GFP-DAD1 and H2a was detected by coimmunoprecipitation (Fig. 2n), which was significantly increased upon proteasomal inhibition, suggesting that molecules targeted for degradation have a higher probability of interacting than newly made ones. In summary, upon proteasomal inhibition DAD1 accumulates in the ERQC, in association with an ERAD substrate and separate from the rough ER, which contains forward translocons.

### OST subunits promote the degradation of glycosylated H2a and to a lesser extent of its non-glycosylated mutant

To investigate a possible role of OST in ERAD, we first overexpressed GFP-DAD1 and analyzed its effect on H2a. Overexpression of GFP-DAD1 led to a dramatic decrease in the levels of H2a (Fig. 3a), suggesting that the protein was degraded. There was no change in the migration of H2a, indicating that glycosylation was not affected. We then partially knocked down DAD1 (Suppl. Fig. 1), which had a major effect in increasing the levels of H2a, suggesting that ERAD was inhibited (Fig. 3b). This partial knockdown had only a very minor effect on glycosylation, as a weak band of faster migration appeared suggesting the existence of a small population of underglycosylated molecules (2 of the 3 glycosylation sites occupied according to our previous studies ^37^). We then tried partially knocking down Tusc3 (Suppl. Fig. 1), which also caused a major accumulation of H2a (Fig. 3c). Tusc3 is a non-essential OST subunit, but nevertheless also caused the appearance of a minor population of underglycosylated H2a molecules. Therefore, partial knockdown of DAD1 or Tusc3, in conditions that affect minimally OST function in glycosylation, have a major effect on inhibiting ERAD.

**Fig. 3.**
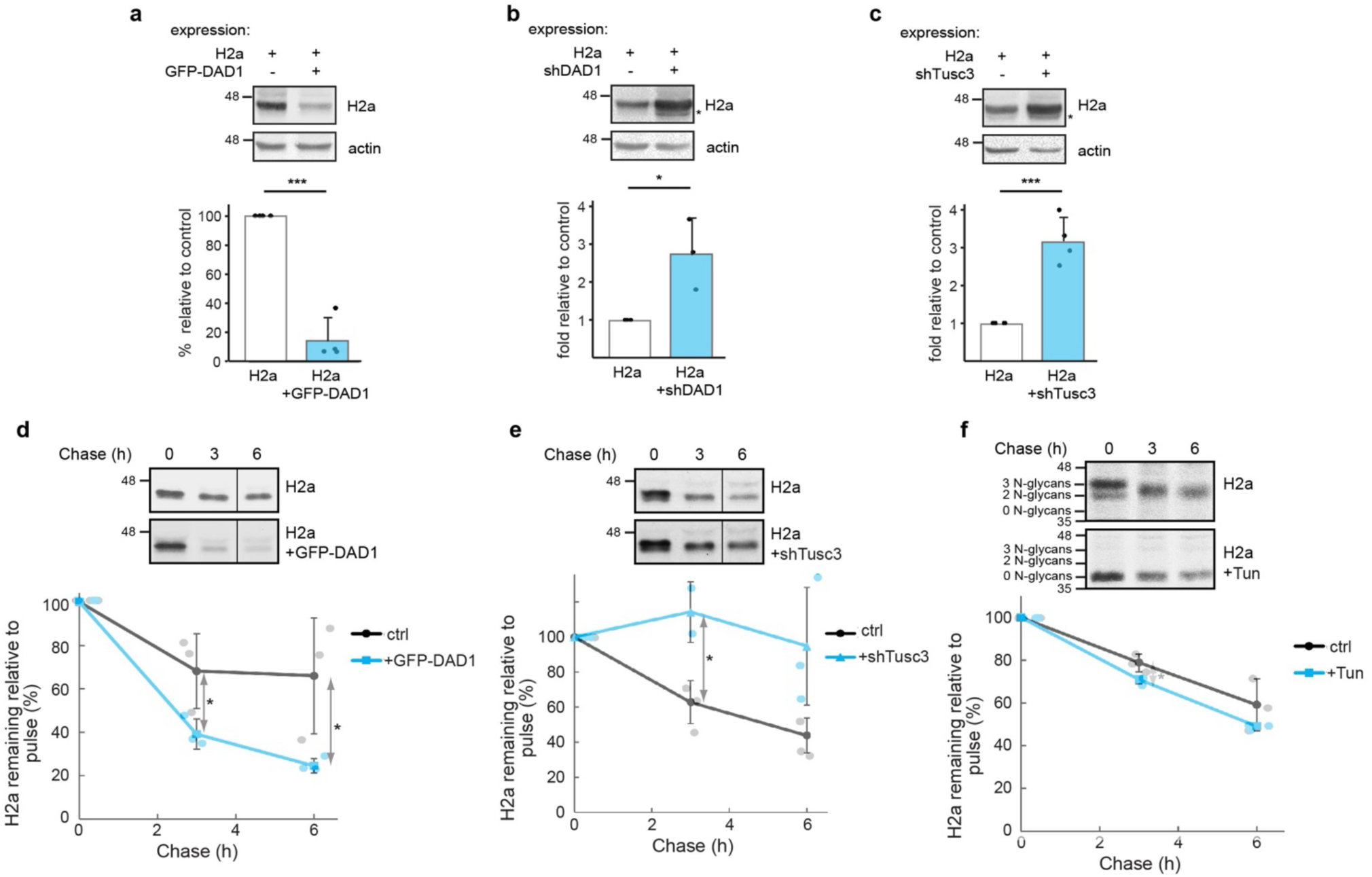
OST subunits promote degradation of a glycosylated ERAD substrate. **(a-c)** Overexpression of GFP-DAD1 strongly reduces the levels of H2a, knockdown of DAD1 or Tusc3 increases H2a levels. Cell lysates were immunoblotted with anti-H2a amino-terminal antibody. The graphs show averages of 3 or 4 independent experiments ±SD. p value ±GFP-DAD1 = 2x10^-5^, ±shDAD1 = 0.03, ±shTusc3 = 0.0004. **(d-f)** Pulse-labeling for 20 min with [^35^S]cysteine and chase for the indicated times of cells expressing H2a with or without overexpression of GFP-DAD1 (d), knockdown of Tusc3 (e) or treatment with Tun (5 μg/ml) for 18h prior to the pulse and during the pulse and chase times (f). H2a degradation (percentage remaining relative to pulse) was accelerated by GFP-DAD1, inhibited by Tusc3 knockdown and was not affected by blocking glycosylation (+Tun). The graphs show averages of 3 independent experiments ±SD. p value ±GFP-DAD1 3h = 0.05, 6h = 0.05; ±shTusc3 3h = 0.03; ±Tun 3h = 0.04.

To confirm that the effects on the steady state levels of H2a were due to changes in its degradation, we performed pulse-chase analysis experiments using [^35^S] cys metabolic labeling. The overexpression of GFP-DAD1 led to a significantly accelerated degradation of H2a (Fig. 3d). Conversely, knockdown of Tusc3 strongly slowed down the degradation (Fig. 3e).

Although glycosylation of H2a did not seem to be affected, we wanted to test whether the effect of Tusc3 knockdown on H2a degradation could be due to inhibition of N-glycosylation of key components of quality control or ERAD. To test this, we inhibited completely N-glycosylation by a long (18h) incubation of cells with tunicamycin (Tun), after which we performed a pulse-chase experiment in the presence of Tun. The treatment caused a shift in the migration of H2a due to lack of its 3 N-glycan chains (Fig 3f). However, the rate of H2a degradation showed no significant change in the presence of Tun (Fig 3f graph), which suggests that the effect of Tusc3 knockdown in delaying degradation (Fig. 3c and 3e) cannot be ascribed to inhibition of glycosylation of the ERAD substrate or of ERAD components.

Similar experiments were performed on a non-glycosylated mutant of H2a (H2aΔgly, ^37^). GFP-DAD1 overexpression and Tusc3 knockdown had similar effects on H2aΔgly as on glycosylated H2a, but to a much lesser extent (compare Fig. 4 a, c, d with Fig. 3 a, c, e respectively). DAD1 knockdown did not have any effect on H2aΔgly degradation (Fig. 4b). DAD1 does not seem to participate in H2aΔgly ERAD at its endogenous levels, but can target the non-glycosylated protein when DAD1 is overexpressed.

**Fig. 4.**
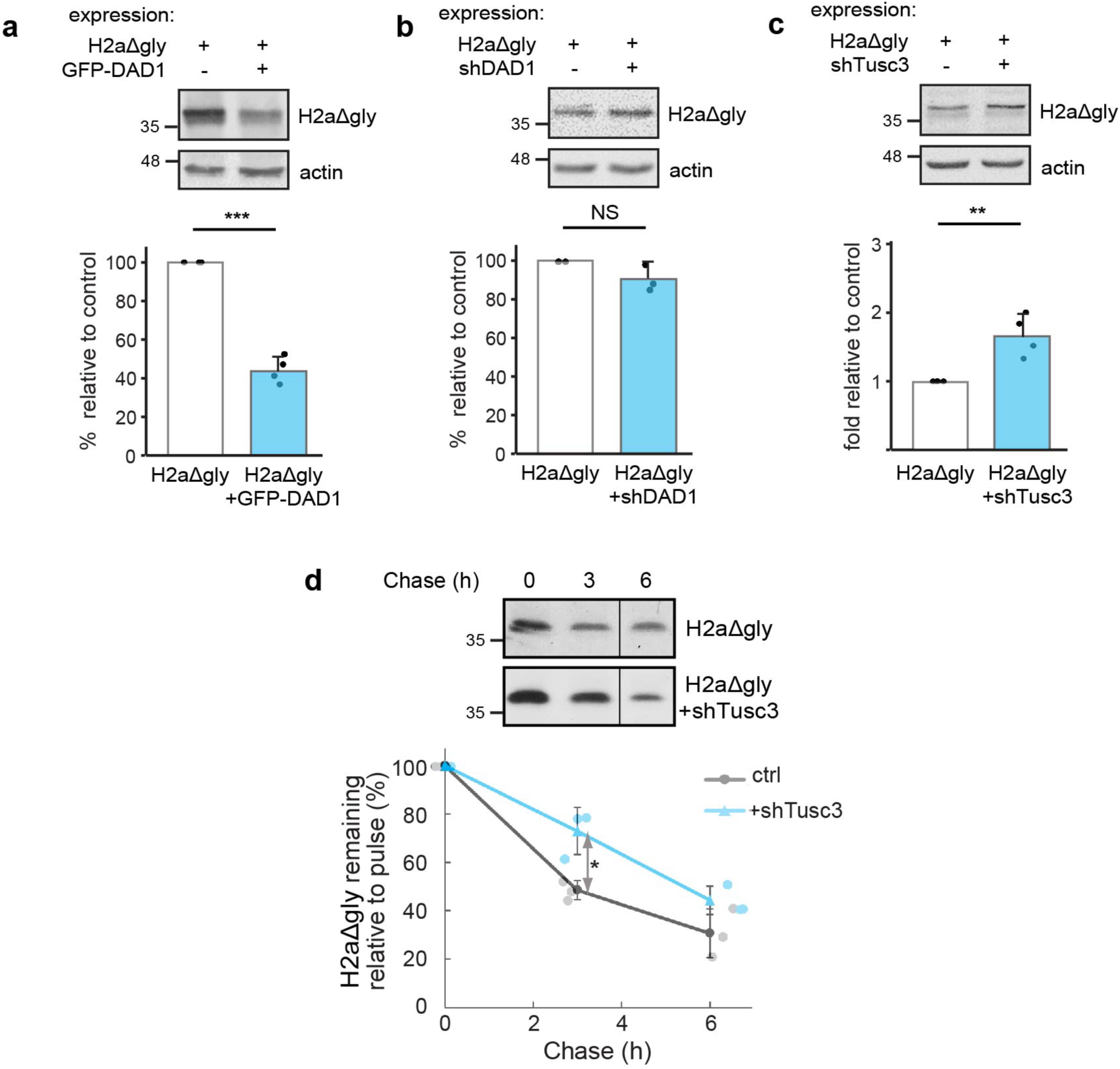
Participation of OST subunits to a lesser extent in the degradation of an unglycosylated ERAD substrate. **(a-c)** Overexpression of GFP-DAD1 reduces the levels of H2aDgly, knockdown of DAD1 does not affect its levels whereas Tusc3 knockdown increases moderately H2aDgly levels. Cell lysates were immunoblotted with anti-H2a amino-terminal antibody. The graphs show averages of 3 or 4 independent experiments ±SD. p value ±GFP-DAD1 = 3x10^-6^, ±shTusc3 = 0.004. **(d)** Pulse/ chase with [^35^S]cysteine of cells expressing H2aDgly shows a moderate inhibition of its degradation by Tusc3 knockdown. The graph shows an average of 3 independent experiments ±SD. p value ±shTusc3 3h = 0.01.

### OST subunits participate in ERAD of other membrane and luminal substrates

To test whether OST subunits are more generally involved in ERAD, we analyzed other well-established ERAD substrates, BACE476, a mutant type I membrane-bound glycoprotein (in contrast with H2a, which is type II, ^11^) and the soluble luminal misfolded glycoprotein human α1-antitrypsin variant null Hong Kong (NHK) ^38^. Overexpression of GFP-DAD1 led to a decrease in the levels of BACE476 and NHK, although to a much lesser extent than for H2a (compare Fig. 5 a, d to Fig. 3a). Similarly, DAD1 or Tusc3 knockdown caused an accumulation of BACE476 and NHK but again less pronounced than for H2a (compare Fig. 5 b, c, e, f to Fig. 3 b,c).

**Fig. 5.**
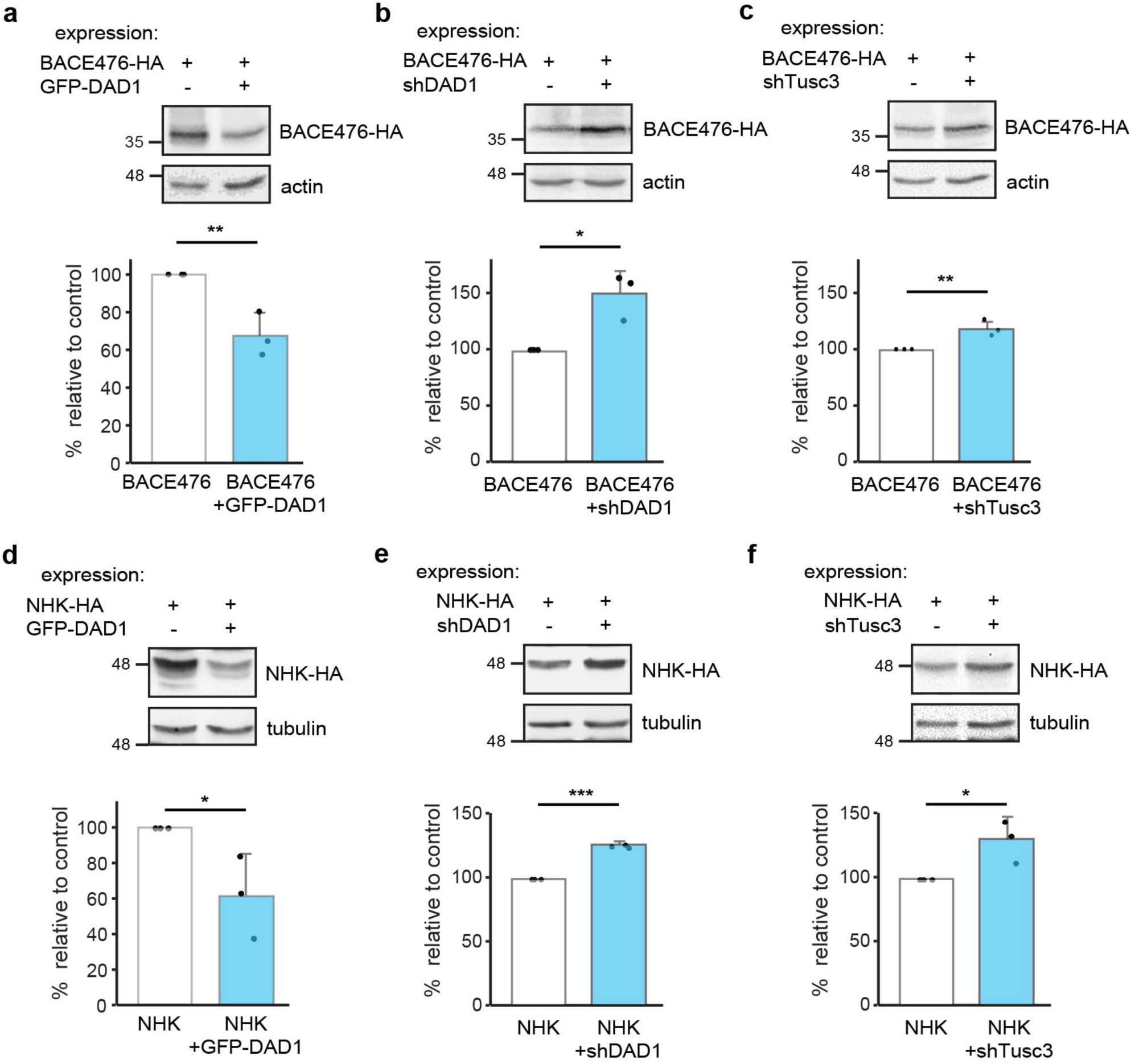
OST subunits affect levels of other ERAD substrates. **(a-c)** Overexpression of GFP-DAD1 reduces moderately the levels of BACE476-HA, knockdown of DAD1 or Tusc3 cause a minor increase in BACE476-HA levels. Cell lysates were immunoblotted with anti-HA and anti-actin as loading control. **(d-f)** Overexpression of GFP-DAD1 causes a minor reduction in the levels of NHK-HA, knockdown of DAD1 or Tusc3 increase moderately NHK-HA levels. Cell lysates were immunoblotted with anti-HA and anti-β tubulin as loading control. The graphs show averages of 3 independent experiments ±SD. p value for BACE476-HA: ±GFP-DAD1 = 0.008, ±shDAD1 = 0.01, ±shTusc3 = 0.003; for NHK-HA: ±GFP-DAD1 = 0.05, ±shDAD1 = 2x10^-6^, ±shTusc3 = 0.03.

Thus, OST subunits can promote ERAD of type I or type II transmembrane substrates and of a soluble luminal substrate, but with some differences in the extent of their action depending on the substrate.

### Interaction between DAD1 and HRD1 and promotion of retrotranslocation by OST subunits

As the ERAD substrates used in our experiments are targeted to HRD1, we wondered whether DAD1 and HRD1 interact. In a coimmunoprecipitation experiment, we could see a robust and specific interaction between GFP-DAD1 and HRD1-myc (Fig. 6a). Although both DAD1 and HRD1 accumulated upon proteasomal inhibition, the ratio of DAD1 coimmunoprecipitated in association with HRD1 decreased (Fig. 6a graph). This suggests that possibly DAD1 associates with HRD1 in a functional ER membrane complex and does not interact with HRD1 molecules that are already targeted for proteasomal degradation.

**Fig. 6.**
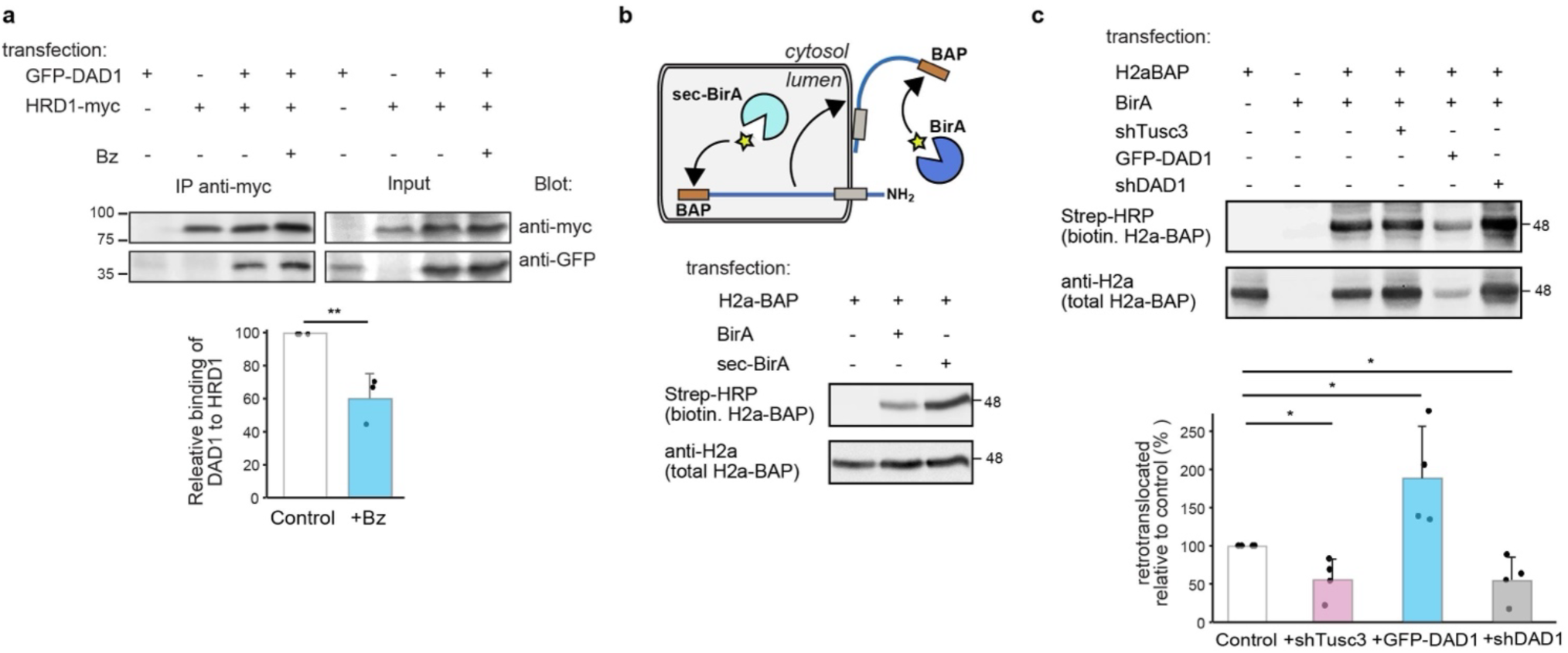
Interaction of an OST subunit with HRD1 and promotion of retrotranslocation of an ERAD substrate. **(a)** Immunoprecipitation with anti-myc of lysates of HEK293 cells expressing HRD1-myc together with GFP-DAD1 and treated with or without 2 μM Bz for 4 h. Immunoprecipitates and 10% of the total lysates (input) were immunoblotted with anti-myc and anti-GFP. The graph shows reduced coimmunoprecipitation of GFP-DAD1 with HRD1-myc under proteasomal inhibition. Values are relative to control (=100), normalized by total GFP-DAD1 and HRD1-myc, average of 3 independent experiments ±SD. p value = 0.009. **(b)** Scheme of addition of biotin (star) to H2a-BAP by an ER luminal biotin ligase, sec-BirA or after retrotranslocation by a cytosolic one, BirA. HEK293 cells transiently expressing H2a-BAP with or without BirA or Sec-BirA were incubated 24h post-transfection with biotin (100μM) for 30 min before lysis. Strep-HRP was used to detect biotinylated H2a-BAP and anti-V5 to detect total H2a-BAP. **(c)** Cells expressing H2a-BAP together with BirA and GFP-DAD1 overexpression or knockdown of DAD1 or Tusc3 were incubated with biotin for 3h. Cell lysates were reacted with Strep-HRP or with anti-H2a antibody. Retrotranslocated (biotinylated) H2a-BAP increased almost two-fold under GFP-DAD1 overexpression and was reduced to half under Tusc3 or DAD1 knockdown. Values are averages of 4 independent experiments ±SD, relative to total H2a-BAP and standardized by the control (=100). P value +shTusc3 vs. control = 0.02, +GFP-DAD1 vs. control = 0.04, +shDAD1 vs. control = 0.02.

We next studied the requirement of OST subunits for the retrotranslocation step. We tested the effect of manipulating the levels of OST subunits on retrotranslocation using the BAP/BirA system ^39^. This system utilizes the E. coli biotin ligase BirA expressed in the cytosol of mammalian cells to specifically biotinylate a 15 amino acid biotin-acceptor peptide (BAP). Secretory proteins with a BAP tag on their ER luminal side are biotinylated only after retrotranslocation. A construct of H2a with BAP attached to its luminal C-terminus behaved consistently as expected for a type II membrane protein ^40^. H2a-BAP was biotinylated poorly when coexpressed with cytosolic BirA, but to a much higher extent when coexpressed with a luminally expressed secBirA, which contains a signal peptide for translocation into the ER ^39^ (Fig. 6b). There was no biotinylation in a control without BirA or secBirA. As BAP is on the luminal C-terminus of H2a, it can be exposed to the cytosolic BirA only after H2a-BAP retrotranslocation. Overexposure of the cytosolic biotinylated retrotranslocated H2a signal allowed us to assess the effect of overexpression or knockdown of OST subunits (Fig. 6c). GFP-DAD1 overexpression caused a decrease in total H2aBAP, as observed for H2a (Fig. 6c lower panel, compared to Fig. 3a), but there was a much smaller decrease in retrotranslocated biotinylated H2aBAP (Fig. 6c upper panel), resulting in a large increase in the ratio of retrotranslocated molecules relative to total, as compared to the control (Fig. 6c graph). Knockdown of either DAD1 or Tusc3 had the opposite effect, resulting in a significant decrease in the retrotranslocation ratio compared to the control.

Together, these results suggest that OST interacts with HRD1 and participates in the retrotranslocation process towards ERAD.

### Molecular dynamics simulation suggest membrane thinning by OST

The molecular architecture of the human OST complexes had been determined by cryo-EM ^41^, showing a dense region traversing the ER membrane with multiple transmembrane domains. (Fig. 7 a,b). Molecular dynamics simulations (pre-rendered at the MemProtMD database ^42^) indicate substantial membrane thinning by the OST transmembrane domains (Fig. 7 c,d). This may lower the energy barrier for protein retrotranslocation, or dislocation from the membrane, as proposed for the interphase between HRD1 and Der1 in yeast ^43^.

**Fig. 7.**
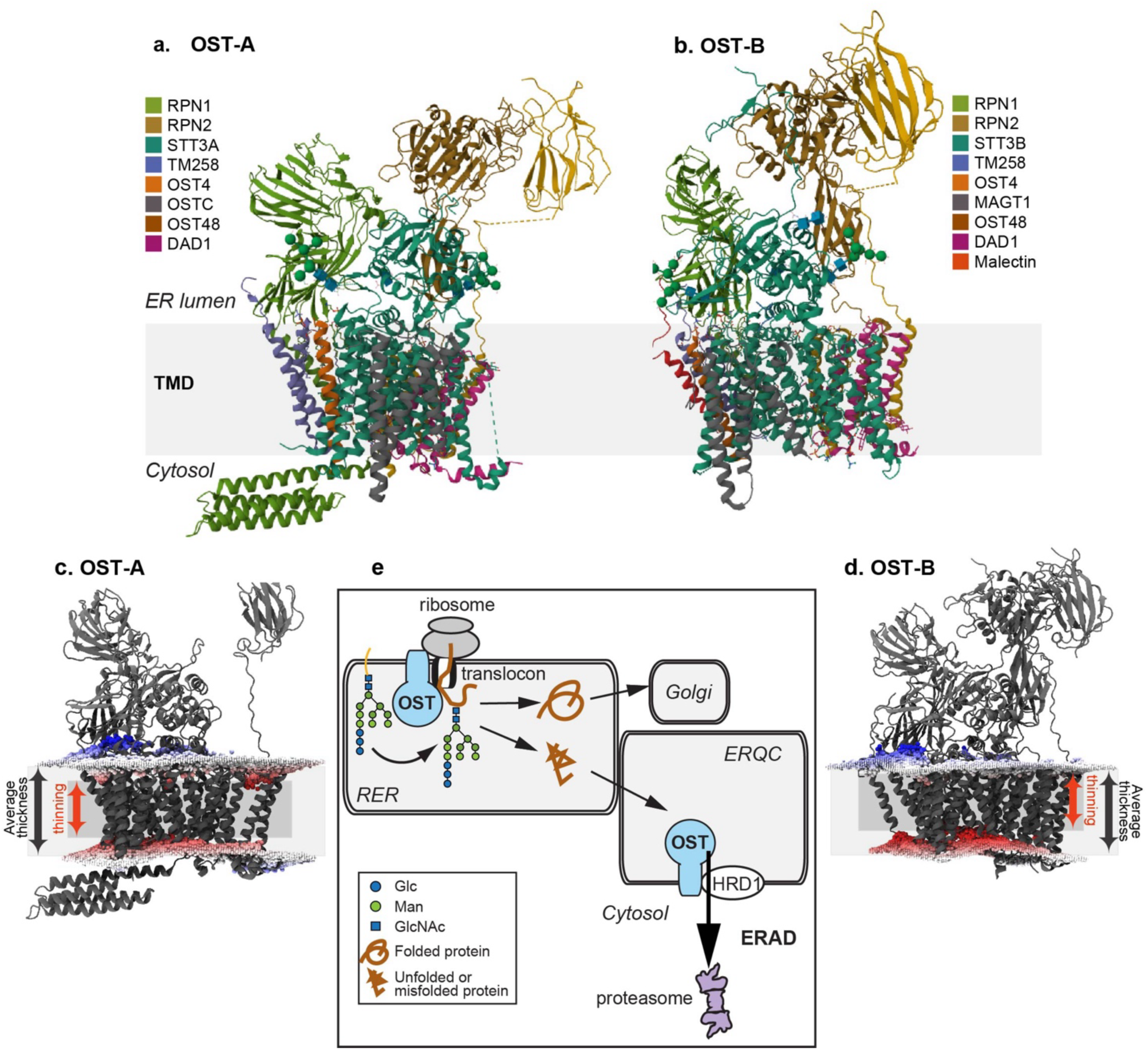
Membrane thinning by OST complexes. **(a,b)** 3D structures of human oligosaccharyltransferase complexes OST-A and OST-B determined by cryo-EM at 3.5 Å resolution, indicating with different colors the observed subunits and illustrating the topology in the ER in relation to the transmembrane domain (TMD). From the RCSB database (6S7O and 6S7T). **(c, d)** Side views of OST-A and OST-B with membrane boundaries from simulations by molecular dynamics pre-rendered at the MemProtMD database. Distortions of the lipid bilayer indicate considerable membrane thinning by the OST transmembrane domains, highlighted by red lipid phosphate head groups, whereas blue illustrates thickening. **(e)** Model illustrating the interaction of OST with translocon components during forward translocation and its participation in N-glycosylation. OST also interacts with HRD1 and participates in retrotranslocation and targeting to ERAD.

## Discussion

In the present study we aimed to identify ERAD machinery components through their interaction with a model misfolded protein being targeted to this pathway. For this aim, we developed an approach for isolation of functional protein complexes engaged in the retrotranslocation event, under conditions in which there is enrichment in retrotranslocation complexes, while minimizing the abundance of forward translocons. This approach employed proteasomal inhibition to accumulate the ERAD substrate just prior to degradation. Our previous studies with H2a, the model substrate used, indicated that in these conditions H2a accumulates in the ERQC membrane, because its retrotranslocation appears to be coupled to its degradation ^11,12,24,32^. To reduce the presence of components of the forward translocon in the protein complexes obtained, we incubated the cells with puromycin. This antibiotic causes release of nascent chains from the ribosome and as a result from the translocons. The combination of puromycin treatment with the C-terminal SBP tag used as bait was intended to prevent isolation of the machineries in charge of translation and forward translocation through incompletely translated H2aSBP molecules (which would lack a complete SBP). In these conditions, we observed an association of several of the OST subunits along with H2a in pulldown experiments. Notably, some of these subunits, OSTC, STT3A and MAGT1 were detected only after proteasome inhibition (Fig. 1g); this, together with the puromycin treatment suggests a preferential interaction with the ERAD substrate after its translation and translocation. Consistently, coimmunoprecipitation of DAD1 occurred after full glycosylation of H2a (Fig. 2n).

OST association was reported with other ER proteins in different high MW complexes, one of them containing SSRD (Trap delta) ^44^. Interestingly, we also detected SSRD only after puromycin treatment and proteasome inhibition (Fig. 1g). Mutations in SSRD lead to defects in glycosylation ^45^, suggesting an early cotranslational event, as the SSR/ Trap complex has been implicated in forward translocation. However, an alternative role has also been suggested for this complex in ERAD ^46,47^. Other ER proteins have been suggested to participate in both forward translocation and in retrotranslocation towards ERAD, such as calumin (CCDC47) ^48^, and the Sec61 complex, for which interaction was found with ERAD factors ^34^. Consistently, we found Sec61 subunits and CCDC47 at similar levels in cells with and without proteasomal inhibition (Fig. 1g). It is noteworthy that TDP-43 appeared enriched in cells with proteasomal inhibition. TDP-43 is localized mainly in the nucleus but has been found sequestered at the cytosolic face of the ER in ALS ^49^.

For OST, noncanonical activities have also been suggested, such as that required for Dengue virus infection, which could be sustained with catalytically inactive OST subunits ^50^. The OST subunit MAGT1 and its oxidoreductase activity ^51^ appeared essential for the infection. MAGT1 was present as one of the most significant interactors of our substrate under proteasome inhibition (Fig. 1g). MAGT1 is found associated with an STT3B containing complex ^52^. The presence of MAGT1 and the effects of knockdown of Tusc3 (also associated with STT3B) would implicate the STT3B complex in our study. STT3B was not detected in the proteomics results, perhaps due to low abundancy, but STT3A and its associated OSTC were, suggesting also a participation of the STT3A complex in ERAD (Fig. 1g).

There are previous studies that hint to the participation of OST subunits in quality control and ERAD. It was suggested that ribophorin I might be involved in protein quality control, as it interacted with a quality control substrate, a fragment of the amyloid precursor protein, after its translocation, remaining associated after puromycin release ^21^. In the case of ERAD highjacking by CMV, the CMV US11 protein was found associated with OST subunits, ribophorin 1, ribophorin 2, and OST48^53^. The ER protein malectin forms a complex with ribophorin 1 which retains misfolded NHK α1-antitrypsin and misfolded ribonuclease A but not the WT counterparts in the ER ^20^. In a proteomics study using OS-9 as a bait, Christianson et al pulled down ribophorin 1 with high score ^54^. They also pulled down other OST subunits, such as STT3A, STT3B and DAD1 using UbxD8, Derlin-1 or Derlin-2 as baits ^15^. However, these observations were not further investigated.

Our proteomics findings, along with insights from the effects of modulating DAD1 and Tusc3 levels, strongly suggest the participation of most, if not all, OST subunits in ERAD. It is likely that DAD1 and Tusc3 fell below the detection threshold in our proteomics analysis. Despite this, DAD1 exhibited clear associations with the ERAD substrate (Fig. 2n) and HRD1 (Fig. 6a) and showed strong effects of its overexpression. The effect of overexpression of this single OST subunit on ERAD might be explained by the limited quantity of its endogenous expression in relation to other subunits ^55^. Interestingly, both DAD1 and Tusc3 partial knockdowns significantly affected ERAD without any significant change in glycosylation.

Both DAD1 and Sec61β relocalized to the ERQC under proteasomal inhibition (Fig. 2), suggesting that most of their molecules are associated with the ERQC and not with forward translocons in these conditions. In a previous study, GFP-DAD1 had been observed to remain with relatively low mobility in the ER membrane, even after puromycin termination of protein synthesis and release from translocons. This observation led to the conclusion that GFP-DAD1 probably remains associated with inactive translocons ^31^. However, the low mobility after puromycin treatment (as we do here) might instead signify a population of GFP-DAD1 molecules associated with a retrotranslocation complex. Indeed, GFP-DAD1 associates with HRD1 (Fig. 6a). Intriguingly, other OST subunits were reported to undergo cleavage by the intramembrane rhomboid protease RHBDL4 before being targeted to ERAD ^56^. Thus, if GFP-DAD1 were undergoing a similar targeting for ERAD, we would expect to observe cleaved molecules upon association with HRD1, but this was not the case.

In conclusion, OST appears to have a dual role, of N-glycosylation at the initial stages of protein translocation into the ER and aiding ERAD at the final stages in the life of misfolded molecules (Fig. 7e). We can only speculate, with the data that we have at the moment, about how OST precisely contributes to ERAD. The fact that the OST complex encompasses many transmembrane domains, interacts with HRD1, affects retrotranslocation and is predicted to cause membrane thinning suggests that it may participate directly in the mechanism of retrotranslocation. It might contribute to membrane distortion, directly aiding retrotranslocation, similarly to the mechanism proposed for the interphase between HRD1 and Der1 in yeast ^43^.

Targeting to ERAD has been best studied for glycoproteins and is linked to glycan processing and recognition steps ^12^. OST could be a candidate for glycan recognition and targeting to ERAD. However, OST function for ERAD does not appear to be restricted to glycoproteins, although the results of OST subunit overexpression and knockdown suggest some preference for glycosylated substrates (Fig. 3, 4). This dual targeting could be equated to the activities of OS-9, XTP3-B and EDEM1 in targeting both glycoproteins and non-glycoproteins to ERAD ^37,57,58^.

## Materials and Methods

### Materials

Streptavidin-agarose-beads, Biotin, tunicamycin, puromycin, MG-132, ALLN, bortezomib, digitonin were from Sigma. Lactacystin was from Cayman Chemicals (Ann Arbor, MI). Promix cell labeling mix ([35S]Met plus [35S]Cys), 1000 Ci/mmol, was from PerkinElmer Life Sciences. Predesigned shRNAs for Tusc3 (TRCN0000038065, TRCN0000038067, TRCN0000299038, TRCN0000299109, TRCN0000299107) and DAD1 (TRCN0000310240, TRCN0000200464, TRCN0000083039, TRCN0000200404, TRCN0000083041) were from Sigma.

### Antibodies

Rabbit anti-DAD1, anti-Tusc3 and anti-calnexin, and mouse anti-β tubulin and anti-actin were from Sigma. Mouse anti-HA was from Biolegend and anti-Myc was custom produced and used before ^35^. Mouse anti-rRNA from Abcam was a kind gift of Orna Elroy-Stein. Rabbit anti-RPL26 was from Cell Signaling. Rabbit polyclonal anti-H2a carboxy-terminal and amino-terminal antibodies and mouse monoclonal anti–H2a carboxy-terminal antibody B9 were the ones used in earlier studies ^11,59^. Mouse anti–S-tag was from Novagen (Gibbstown, NJ). Rabbit anti-GFP was from Santa Cruz Biotech. Rabbit anti-RFP was from MBL. Rabbit anti-Sec61β was the one used previously ^32^. Strep-HRP, goat anti-mouse IgG-HRP, goat anti-rabbit IgG-HRP and goat anti-mouse IgG conjugated to Cy2 or to Dylight 549 were from Jackson-Immuno-Research Labs.

### Plasmids and constructs

WT HRD1-myc, dominant negative mutant hsHRD1-myc (MutHRD1-myc), Sec61β-GFP, S-tagged-OS-9.1 and S-tagged-OS-9.2 in pcDNA3.1(+), H2a and H2a mutated in its three glycosylation sites (H2aΔgly), subcloned in pCDNA1 and BACE476-HA were used before ^11,24,36,60^, as well as human α1-antitrypsin NHK-HA in pcDNA3.1 ^27^. Constructs encoding H2aG78R uncleavable mutant fused through its C-terminus to a 38-amino acid streptavidin binding peptide (H2aSBP), to monomeric red fluorescent protein (H2aRFP) or to GFP (H2aGFP) were described before ^11,23,24^. DAD1 in pEGFP-C1 (GFP-DAD1) was a kind gift from Gert Kreibich. H2a-BAP-V5 was used before ^40^. BirA and Sec-BirA plasmids were a kind gift of Gianlucca Petris and Oscar Burrone (ICGEB, Trieste, Italy) ^39^.

### Cell culture, media and transfections

NIH 3T3 and HEK293 cells were grown in DMEM supplemented with 10% bovine calf serum at 37 °C under 5% CO_2_. Transfections of NIH 3T3 cells were carried out using a Neon MP-100 microporator system (Life Technologies, Carlsbad, CA). Transfection of HEK293 cells was performed using the calcium phosphate method. An HEK293 stable cell line expressing H2aSBP (SBP6) was described previously ^23^.

### BAP construct transfection and probing

H2a-BAP-V5 was transfected using the calcium phosphate method in HEK293 cells together with an equal amount of plasmid carrying BirA (cytosolic biotin ligase). Biotin (100μM) was added 24 hours post transfection overnight. The cells were then rinsed with PBS and lysed at 4°C in Buffer A (1% Triton X-100, 0.5% NaDOC and 1X cOmplete Protease Inhibitor Cocktail (Roche) in PBS), containing 10mM N-ethylmaleimide, to block BirA post-lysis activity. The probing of biotinylated proteins on blots was performed using Strep-HRP in PBS-TWEEN (0.5%), followed with detection by enhanced chemiluminescence (ECL).

### Metabolic labeling

Subconfluent (90%) cell monolayers in 60-mm dishes were labeled for 30 min with [^35^S]Cys and chased for different periods of time with normal DMEM plus 10% FCS, lysed, and immunoprecipitated with anti-H2a carboxy-terminal antibody as described previously ^59,60^. Quantitation was performed in a Fujifilm FLA 5100 phosphorimager (Tokyo, Japan).

### Immunoprecipitation, crosslinking and immunoblotting

Cell lysis in buffer A and immunoprecipitation methods are described in ^23^. For GFP-DAD1 co-expressed with H2a, crosslinking was performed with DSP prior to cell lysis. Cells were washed 2X with PBS and incubated with PBS containing 1mM DSP at 4°C for 1 hour. The reaction was stopped by incubation at 4°C for 20 min with 50mM Glycine, followed by a rinse with PBS and incubation with 20mM Tris-HCl pH7.4 in PBS for at RT 5 min.

For precipitation with streptavidin-agarose beads, cell lysates were incubated with the beads on a vertical rotating mixer at 4°C overnight. After centrifugation, the beads were washed three times with 0.5% Triton X-100, 0.25% NaDOC and 0.5% SDS in PBS and once with PBS. Samples were boiled in sample buffer for 5 minutes and run on SDS-PAGE under reducing conditions.

Transfer to a nitrocellulose membrane, immunoblotting, detection by ECL, and quantitation in a Bio-Rad ChemiDocXRS Imaging System (Hercules, CA) were done as described previously ^35^. Quantity one or ImageJ software were used for the quantitation.

### Iodixanol equilibrium sedimentation gradient

Iodixanol gradients were performed similarly to previously described ^27^. Briefly, HEK293 cells expressing H2a-RFP (untreated or treated with Bz) were washed with PBS, resuspended in a homogenization buffer (0.25M Sucrose, and 10mM HEPES pH7.4). The cells were passed through a 21G needle 5 times before homogenization in a Dounce homogenizer (low clearance pestle, 30 strokes). The homogenates were centrifuged at 1000xg for 10 minutes at 4°C to remove nuclei and cell debris and the supernatants were loaded on top of an iodixanol gradient (10 to 34%). The gradients were ultra-centrifuged at 24,000 rpm (98,500g, Beckman SW41 rotor) at 4°C for 16 hours. 1ml gradient fractions were collected from top to bottom, precipitated with 10% TCA and run on SDS-PAGE.

### Immunofluorescence Microscopy

Immunofluorescence was performed as described previously ^32,61,62^. Briefly, cells grown for 24 h after transfection on coverslips in 24-well plates were fixed with 3% paraformaldehyde for 30 min, incubated with 50mM glycine in PBS, and permeabilized with 0.5% Triton X-100. After blocking with normal goat IgG in PBS/ 2% BSA, they were exposed to primary antibody for 60 min, washed and incubated for 30 min with a secondary antibody, followed by washes and staining of nuclei with DAPI. Specimens were observed on a Zeiss laser scanning confocal microscope (LSM 510; Carl Zeiss, Jena, Germany). ImageJ was used to quantify fluorescence intensity and to calculate Mander’s coefficients (using JACOP) for colocalization studies. ERQC localization was determined by the accumulation of H2a fluorescent fusion proteins or of OS-9.

### Proteomics

SBP binding and elution conditions were as in ^63^). For SILAC proteomics HEK293 or SBP6 cells were labeled with 13C(6)15N(2) L-lysine for 10 days. The labeling efficiency obtained was 97%. 7x10^8^ cells (15x150mm dishes) were treated with or without 50μM MG-132 + 50μM ALLN for 4h and with 2mM puromycin for the last 15min before lysis. Cells were lysed in 1% digitonin in PBS+2mM PMSF+5μg/ml aprotinin. Labeled HEK293 and unlabeled SBP6 lysates were mixed at a 1:1 protein ratio. The combined lysates were subjected to ON precipitation with preblocked streptavidin beads (0.5% BSA, 0.2% digitonin in PBS for 2-3 h, washed once with 0.1% BSA in the same buffer composition). Precipitation was with beads in 0.1% BSA, 0.2% digitonin in PBS, followed by 4 washes with 0.2% digitonin in PBS and protein interactors pulled down with H2aSBP were eluted for 1h at +4°C by competition with 250μM free biotin dissolved in the wash buffer. Samples were heated at 65°C with β-mercapto-ethanol containing sample buffer for 10min and run on SDS-PAGE. Proteins were identified by nano LC-MS/MS using a nanoACQUITY UPLC nano liquid chromatography system (Waters) coupled to a LTQ orbitrap mass spectrometer (Thermo) or an EasyLC system (Proxeon) coupled to a Thermo LTQ-FT Ultra mass spectrometer. For nanoLC peptide separation, 15 cm long capillary column (75 μm inner diameter) with a prepulled capillary tip (Proxeon) were packed in-house with Reprosil-Pur 3 μm C18 material (Dr. Maisch). Peptides were eluted at 0.3 μl/min using stepwise gradients from 2%–60% acetonitrile, in 0.1% formic acid. MS/MS analysis on the LTQ orbitrap mass spectrometer was performed with standard settings using cycles of 1 high resolution MS scan from m/z 370-1750 followed by MS/MS scans of the 10 most intense ions with charge states of 2 or higher. For analysis on the Thermo LTQ-FT Ultra mass spectrometer, FT scans from m/z 400-2000 were taken at 100,000 resolution, followed by collision induced dissociation (CID) scans in the LTQ of the 8 most intense ions with signal greater than 2000 counts, and charge state of 2 or higher.

### Bioinformatics analysis

Protein identification and SILAC based quantitation was performed with MaxQuant (version 1.0.13.13) using default settings. The UNIPROT human database (version 2009-04-14) was used for protein identification. MaxQuant uses a decoy version of the specified UNIPROT database to adjust the false discovery rates for proteins and peptides to below 1%.

Gene ontology enrichment of 86 differentially expressed proteins was performed using DAVID tool^64^. Selected enriched functions were presented. <u>D</u>ifferentially expressed proteins were applied to STRING protein-interaction tool ^65^. Protein colors represent enriched functions circled in colors.

### Statistical Analysis

The results are expressed as average ± SD. Student’s t-test (two-tailed) was used to compare the averages of two groups. Statistical significance was determined at P<0.05 (*), P<0.01 (**), P<0.001 (***).

## Acknowledgments

We would like to thank G. Kreibich, O. Elroy-Stein, G. Petris and O. Burrone for plasmids and reagents. We are very grateful to F.U. Hartl for ongoing advice and for critically reading and helping in the edition of this manuscript. Work was supported by grant 2577/20 from the Israel Science Foundation (GZL).

## Author Contributions

Conceptualization: GZL. Investigation: MS, NO-S, CP, BG, S.S., RK. Bioinformatics: MPC. Supervision: GZL. Writing: GZL. Review and editing: GZL, MS, MPC, RK.

## Declaration of Interests

The authors declare no competing interests.

## Notes

### Competing Interest Statement

The authors have declared no competing interest.

### Summary of Updates

Addition of molecular dynamics simulations suggesting membrane thinning by OST

