## Supplementary information for "Oligosaccharyltransferase is involved in targeting to ER-associated degradation"

**Supplementary figures**

Suppl. Fig. 1

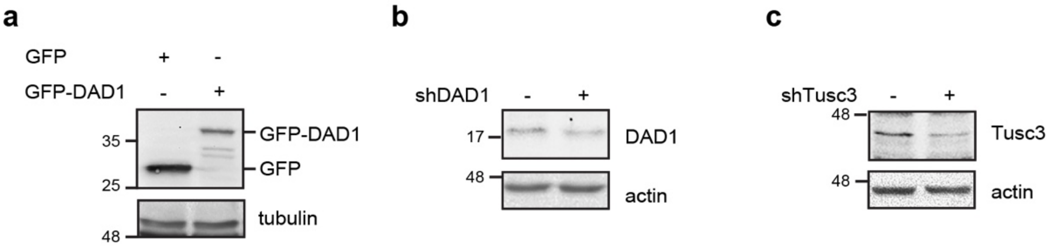


**Suppl. Fig 1. Overexpression and knockdown of OST subunits.** Lysates of HEK293 cells overexpressing GFP-DAD1 or GFP or after knockdown of DAD1 or Tusc3 for 48h, were immunoblotted with anti-GFP (a), DAD1 (b) and Tusc3 (c), with tubulin or actin as loading controls.

**Supplementary Tables**

**Suppl. Table 1.** DAVID function enrichment analysis.

**Suppl. Table 2.** The table shows values for the 86 differentially expressed proteins resulting from the proteomics analysis.

**Suppl. Table 3.** Statistics analysis of the proteomics values.
